# Humanized APP is the primary determinant of regional brain volume in humanized APOE knock-in mice: a cross-sectional ex vivo MRI study

**DOI:** 10.64898/2026.05.04.722764

**Authors:** Avnish Bhattrai, Adam C. Raikes, Roberta Diaz Brinton

## Abstract

**INTRODUCTION:** Age, Apolipoprotein E4 (*APOE*4) genotype, and biological sex are major risk factors for late-onset Alzheimer’s disease (LOAD), and preclinical mouse models enable controlled investigation of these factors. To date, humanized *APOE*4 has not recapitulated LOAD-relevant brain volume phenotypes. Given the central role of amyloid precursor protein (APP) in LOAD pathogenesis, incorporating humanized APP (*h*APP) alongside humanized *APOE* (h*APOE*) may therefore improve translational modeling of structural brain changes characterized by neuroimaging.

**METHODS:** Aged mice (mean age = 23.25 months) carrying murine (*m*) or humanized (*h*) APP and either murine Apoe or h*APOE*3/3 (h*APOE*3-HOM), h*APOE*3/4 (h*APOE*4-HET), or h*APOE*4/4 (h*APOE*4-HOM) underwent in-skull ex vivo volumetric MRI. Regional volumes were quantified in absolute terms and relative to total brain volume (TBV). Linear models included APP type, *APOE* genotype, and sex, with FDR correction applied within contrasts.

**RESULTS:** Brain volumes were primarily determined by APP background, with *h*APP globally reducing total and regional volumes relative to *m*APP mice. Across *h*APP models, h*APOE*4-HOM exhibited the greatest brain-wide reductions, which was mitigated by a single h*APOE*3 allele. In contrast, mouse APP exerted modest effect in h*APOE*, with h*APOE*4 carriers exhibiting greater total volume without regional specificity. After TBV adjustment, *h*APP mice exhibited subcortical vulnerability with relative cortical preservation. Females exhibited larger brain volumes than males, independent of APP or *APOE* genotype.

**DISCUSSION:** h*APP* induces distinctly smaller brain volumes in this humanized APOE knock-in model, and h*APOE*4 homozygosity amplifies that effect, indicating genotype-dependent susceptibility. Because this study is cross-sectional and lacks histopathological confirmation, volume differences may reflect developmental or constitutive effects rather than neurodegeneration. Humanized APP and APOE are therefore necessary but not sufficient to recapitulate the complete volumetric signature of established LOAD. Longitudinal and histological studies are required to determine whether these differences reflect a prodromal trajectory or a developmental effect.

## 1. INTRODUCTION

Late-onset Alzheimer’s disease (LOAD, ∼95% of all AD cases) is driven by multiple genetic and biological factors, including age, biological sex, and *APOE* genotype. LOAD pathology includes cerebral hypometabolism, global brain shrinkage and regionally-selective atrophy, and decreased white matter integrity [1–3]. Early atrophy is evident on MRI in the hippocampus and entorhinal cortex [4–7]. These volumetric decreases are key hallmarks of the LOAD neuroimaging phenotype.

Transgenic preclinical AD mouse models replicate many hallmark LOAD features, including global and regional atrophy, amyloidosis, and inflammation [8–13]. However, these mouse models rely on combinations of rare aggressive familial AD mutations which do not occur together clinically and do not replicate the time course of LOAD pathology, atrophy, or behavior [14] . This limits translational relevance to LOAD biology and progression.

Females and *APOE*4 carriers have the greatest LOAD risk [15, 16]. The MODEL-AD consortium generated humanized *APOE* (h*APOE*) mice to improve mouse model translational relevance for the APOE risk factor [17]. Relative to h*APOE*3/3 (h*APOE*3-HOM) mice, h*APOE*4/4 (h*APOE*4- HOM) mice exhibit lower APOE protein levels, mitochondrial dysfunction, increased immune activation, demyelination, and endocrine profiles consistent with accelerated aging [18–20]. Despite LOAD-relevant phenotypes, cross-sectional MRI studies report modest increases in total brain volume in h*APOE*4 carriers [21] without LOAD-like atrophy or cognitive deficits [22]. Therefore, additional genetic factors (e.g. amyloid precursor protein ) may be necessary to further increase h*APOE* mouse model translational relevance.

While human APP drives amyloid-β plaque formation, murine APP does not aggregate plaques. Existing mouse models using APP overexpression or familial mutations induce AD-like pathology [23–26], and humanized APP (*h*APP) mice exhibit cognitive impairment and reduced long-term potentiation compared to those with murine APP (*m*APP) [27]. However, the impact of *h*APP on brain volume without familial mutations remains unclear.

Studies investigating the combined effect of humanized APP and *APOE* genotype on brain volumetric phenotypes are lacking. Addressing this gap, we investigated the impact of humanized APP on brain morphology using ex-vivo imaging in aged mice (23-25 months) across sex, h*APOE* genotype (3/3, 3/4, 4/4), and APP background (murine vs. humanized).

## METHODS

### 1.1. ANIMAL USE

All procedures were performed following National Institutes of Health guidelines and approved by the University of Arizona Institutional Animal Care and Use Committee (IACUC). Breeder pairs included *m*APP/h*APOE*3, *m*APP/h*APOE*4, and *h*APP/murine Apoe (mApoe) knock-ins (Jackson Laboratories strains #029018 and #027894 on C57BL/6J background, and #030898 on mixed BL/6J and BL/6NJ background). Animals were bred in-house and maintained in ventilated cages in our animal colony (14-h light/10-h dark, ad libitum food and water). A total of 148 male and female *m*APP/h*APOE*3-HOM, *m*APP/h*APOE*3/4 (h*APOE*4-HET), *m*APP/h*APOE*4-HOM, *h*APP/mApoe, *h*APP/h*APOE*3-HOM, *h*APP/h*APOE*4-HET, and *h*APP/h*APOE*4-HOM mice (23-25 months old) were used for these experiments. Sample sizes for each genotype combination are reported in **Figure 1**.

**Figure 1:**
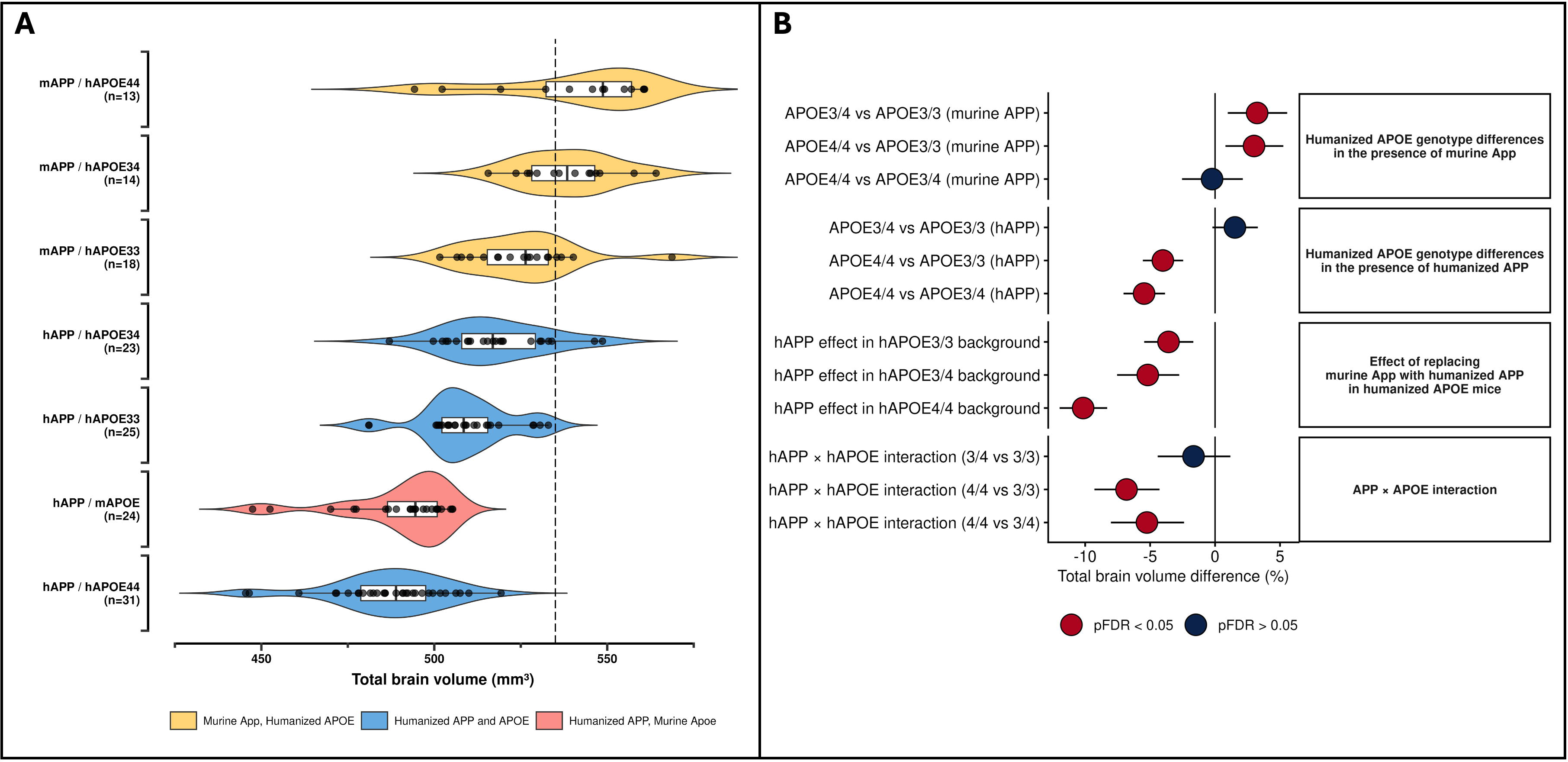
Impact of amyloid precursor protein (APP) type and humanized apolipoprotein E4 (h*APOE*4) carrier status on Total Brain Volume (TBV) showed reduced brain volume in humanized APP carriers relative to their murine counterparts. (A) TBV distribution by APP type and h*APOE*4 carrier status (dashed line represents reference TBV for an adult C57BL/6J mouse), and **(B)** as forest plot showing difference in TBV percentage (%) between groups across different APP types and h*APOE*4 allele carrier combinations. Statistically significant differences in the forest plot (FDR-corrected; p < 0.05) are denoted in red. Error bars represent 95% confidence intervals.

### 1.2. NECROPSY

Mice underwent transcardial perfusion with 1X phosphate buffer saline (PBS) for 4-6 minutes, followed by perfusion with 4% paraformaldehyde (PFA) in PBS for an additional 4-6 minutes. Skulls were cleaned to remove soft tissue and stored in 4% PFA in PBS for 48 hours to allow for complete fixation then transferred to PBS solution with 0.01% sodium-azide for long-term storage.

### 1.3. MRI ACQUISITION

MRI data were acquired on a Bruker Biospec 70/20 7.0T scanner (Bruker Biospin GmbH, Germany) running Paravision 360(v. 3.5). All mouse brains were prepared for ex-vivo imaging by vacuum desiccation in Fluorinert (FC-3283). The ex-vivo imaging protocol included a T2-weighted rapid acquisition with relaxation enhancement (RARE) sequence (TE: 30 ms; TR: 1800 ms; flip angle 180°; RARE factor: 8; number of averages: 2; FOV: 2.4 x 1.44 x 0.96 cm^3^; acquisition matrix 320x192x120; reconstructed voxel size 75μm; total acquisition time: 3 hours and 4 minutes). The *m*APP/h*APOE* mice used here have been previously analyzed by Raikes et al [21], however the analytical pipeline and statistical modeling are different for this study.

### 1.4. DATA PROCESSING

#### 1.4.1. PRE-PROCESSING

Pre-processing steps were carried out as in Raikes et al, 2026 [21]. Briefly, a custom Apptainer container on the University of Arizona’s High-Performance Cluster was utilized for pre-processing and analyzing data. Visual quality control was carried out on T2-weighted images. Preprocessing steps included denoising, intensity scaling, bias field correction (*N4BiasFieldCorrection*; ANTs v. 2.5.4), and creation of native space brain masks using a previously defined template space brain mask [28–31].

#### 1.4.2. ATLAS-BASED WORKFLOW

An initial minimum deformation template (MDT) was generated using a subset of the 148 brains (3 per APP/*APOE*/sex combination), using the *optimized_antsMultivariateTemplateConstruction* pipeline [32]. All brains (n=148) were non-linearly registered to the MDT producing a final study-specific template brain and mask. The template brain was registered to the Allen Mouse Brain template supplied with Transbrain [33]. The Transbrain atlas includes 68 bilaterally labeled regions and was inverse warped to native mouse brain space and summarized as regional. Total brain volume, including brain tissue and ventricles, was defined in native space using the study template mask. Volumetric measurements were recorded in cubic millimeters (mm^3^).

### 1.5. STATISTICAL ANALYSIS

Regional brain volumes were analyzed using linear models fit independently for each anatomical region. Three sets of outcomes were analyzed: total brain volume (TBV), absolute regional volumes, and relative regional volumes adjusted for total brain volume. TBV and all regional volumes were log-transformed prior to analyses.

Animals were grouped according to APP type (murine vs. humanized) and *APOE* genotype (murine Apoe, h*APOE*3-HOM, h*APOE*4-HET, and h*APOE*4-HOM). Therefore, each combination of APP type and *APOE* genotype formed a unique group for the purpose of these analyses (e.g. *h*APP/mApoe, *m*APP/h*APOE*3-HOM, *h*APP/h*APOE*3-HOM, *h*APP/h*APOE*4- HOM).

### 1.5.1. Model specification

Three model families were fit using ordinary least squares in R (v. 4.5.0). Total brain volume was modeled as:

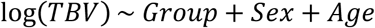

Absolute regional volumes were modeled independently for each region r as:

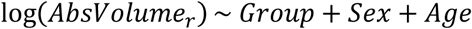

Finally, relative regional volumes were modeled by including log-transformed TBV as a covariate:

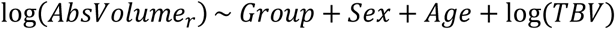

In all models, group denotes APP type/*APOE* genotype groupings. Sex was included as a two-level categorical variable, and age was mean-centered (in days). We could not fit fully parameterized APP x *APOE* interaction models because no mice carried *m*APP/mApoe. For regional analyses, each model was fit once per region and hypothesis tests for group differences were evaluated using estimated marginal means computed using contrast vectors specifying linear combinations of groupings and holding all other variables at their mean. Planned contrasts were used to address biologically motivated comparisons:

1. Differences between h*APOE* genotypes within murine APP (e.g. *m*APP/h*APOE*4- HOM – *m*APP/h*APOE*3-HOM)
2. Differences between h*APOE* genotypes within humanized APP (e.g. *h*APP/h*APOE*4- HOM – *h*APP/h*APOE*3-HOM)
3. The effect of replacing *m*APP with *h*APP within each h*APOE* genotype (e.g. *h*APP/h*APOE*4-HOM – *m*APP/h*APOE*4-HOM)
4. Difference-in-difference contrasts to determine if APP-associated differences differed by *APOE* genotype (e.g. [[*h*APP/h*APOE*4-HOM – *m*APP/h*APOE*4-HOM] – *[h*APP/h*APOE*3-HOM – *m*APP/h*APOE*3-HOM]], indicating whether the *h*APP effect is different in h*APOE*4-HOM mice compared to h*APOE*3-HOM mice).
5. Sex differences independent of APP and *APOE* effects.

For regional analyses, statistical significance was determined using false discovery rate (FDR) correction across regions within each planned contrast. Differences were identified as statistically significant at FDR-corrected p < 0.05.

All volumes were entered into models as log-transformed values. Therefore, groupwise differences were exponentiated and reported as percent differences. Relative regional models include log-transformed TBV as a covariate and therefore group differences are interpreted as percent difference after accounting for overall brain size.

As an exploratory analysis for hypothesis generation, we assessed the relationship between total brain and regional volume with two behavioral indices (novel object recognition discrimination index and open field freezing time, following the protocol used by McLean and colleagues) [22]. These models assessed slope differences in group x volume interactions on behavior. For interpretability, behavioral data were z-scored, and volumes were log-transformed and z-scored prior to analyses and findings are reported at FDR-corrected p < 0.1.

## 2. RESULTS

### 2.1. Total brain volume (TBV)

After adjusting for age and sex, the relationship between *APOE* genotype and total brain volume differed by APP background **(Figure 1A)**. In mice carrying murine APP (*m*APP), the presence of an h*APOE*4 allele was associated with a 3% increase in TBV relative to h*APOE*3-HOM mice (*p*_FDR_ = 0.025-0.031; **Figure 1B)**. Conversely, among humanized APP (*h*APP) mice, h*APOE*4-HOM carriers exhibited 4% (compared to h*APOE*3-HOM; p_FDR_=1.45×10^-5^; d = -1.37 [95% CI: -1.91,-0.83]) to 6% (compared to h*APOE*4-HET; p_FDR_=1.0×10^-8^; d = -1.87 [95% CI: -2.44, -1.31]) smaller brains than those carrying at least one h*APOE*3 allele **(Figure 1B)**.

Comparisons of APP backgrounds within each h*APOE* genotype revealed that humanized APP reduced total brain volume in a genotype-dependent manner. Reductions ranged from 3.6% (d = -1.22 [95% CI: -1.87, -0.56]) in h*APOE*3-HOM mice to 10.15% (d = -3.57 [95% CI: -4.25,-2.89]) in h*APOE*4-HOM mice (p_FDR_ range = 5.6×10^-18^–0.00235; **Figure 1B)**. Difference-in-difference analyses confirmed that the *h*APP-associated reduction was greater in h*APOE*4-HOM background compared to h*APOE*3-HOM (mean difference = 6.8% [95% CI: -9.3%, -4.3%]; d = - 2.35 [95% CI: -3.25, -1.46]; p_FDR_ = 7.1 x 10^-6^; **Figure 1B).**

Ventricular volumes were extracted using the Dorr mouse brain atlas and analyzed using the same linear models. Volumes ranged from 1.20–1.29% of TBV across groups. *h*APP mice had significantly smaller ventricular volumes than *m*APP counterparts (−7.4% to -12.6%, p_FDR_ < 0.001), with the largest reduction in *h*APP/h*APOE*4-HOM mice. After adjustment for total brain volume, these differences were attenuated and non-significant in the *h*APP/h*APOE*4-HOM group (−0.5%, p = 0.81), indicating that ventricles scale proportionally with the globally smaller brain in these animals.

### 2.2. Absolute and TBV-Adjusted Regional Effects of h*APOE* Genotype in mouse APP (*m*APP) Mice

In the *m*APP background, h*APOE*4 carriers exhibited a consistent, brain-wide tendency toward larger volumes relative to h*APOE*3-HOM mice, spanning most cortical and subcortical regions (see **Figure 2A** for region-wise estimates and significance summary and **Figures 2B-2G** for spatial localization of differences). This effect was most evident for h*APOE*4-HET versus h*APOE*3-HOM in a subset of regions, particularly within somatosensory, temporal, hippocampal, striatal, and thalamic areas (p_FDR_ = 0.047–0.049; d = 0.87–1.19 [95% CI: 0.13,1.93]; **Figure 2D**). In contrast, comparisons involving h*APOE*4-HOM and h*APOE*3-HOM did not yield significant spatially localized effects (**Figure 2B-C**), despite trending towards similar volumetric increase as observed for the h*APOE*4-HET carriers.

**Figure 2:**
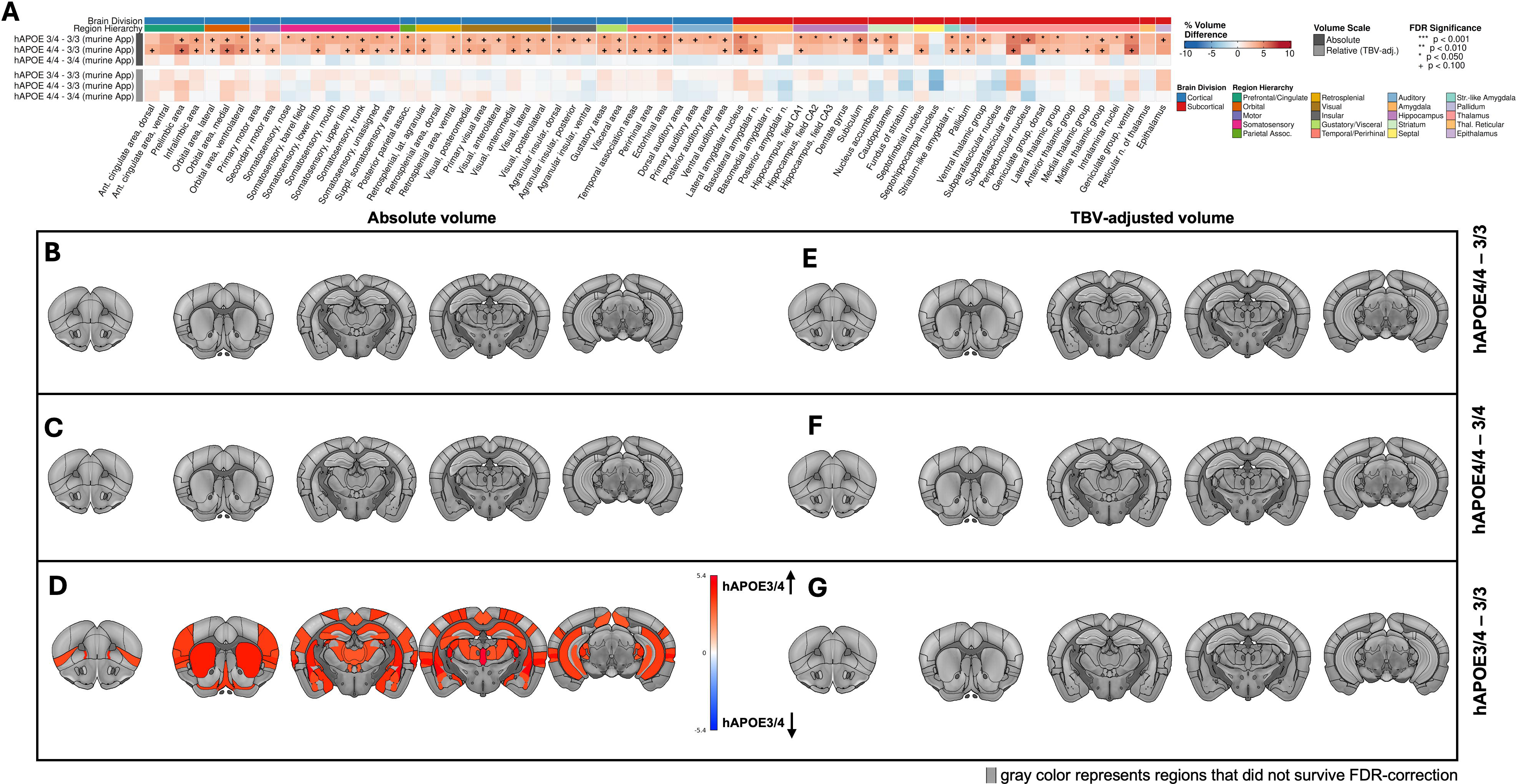
In the presence of murine amyloid precursor protein (APP), each copy of the humanized apolipoprotein E4 (h*APOE*4) allele is linked with increasing total brain volume (TBV). Spatial distribution of regional volume differences (percentage; TBV-adjusted) linked with the effect of carrying at least one copy of h*APOE*4 carrier status relative to h*APOE*3-HOM, on a murine APP background. Greater volume is denoted **in red** and regions with no significant differences **in gray.** All results are FDR-corrected for multiple comparison (p < 0.05). **(A) Cumulative region-by-region difference visualization as a heatmap,** color coded for percentage (%) volume difference, statistical significance and brain region hierarchy. **Left-side** p**anels (B, C and D) represent** absolute (TBV-unadjusted) volume differences while **right-side panels** (**E, F and G)** represent relative (TBV-adjusted) volume differences.

After TBV adjustment, these differences were attenuated (**Figure 2E-G**), with no regional comparisons surviving FDR correction. Together, this diffuse and non-localized profile in h*APOE*4- HOM and HET on the *m*APP background primarily reflects an impact on global scaling of brain volume rather than regionally specific remodeling.

### 2.3. Absolute and TBV-Adjusted Regional Effects of h*APOE* Genotype in *h*APP Mice

In contrast to the *m*APP background, where h*APOE*4 carriers exhibited similar volumetric enlargement compared to the h*APOE*3-HOM mice, the *h*APP background was characterized by similarity between h*APOE*3-HOM and h*APOE*4-HET mice, with reductions observed only in the h*APOE*4-HOM mice (**Figure 3A–D**). In the *h*APP mice, h*APOE*4-HOM was associated with a brain-wide reduction in absolute regional volumes relative to both h*APOE*3-HOM (p_FDR_ = 4.8×10⁻⁷–0.050; d=-0.55 to -2.19 [95% CI: -2.19, -0.01]) and h*APOE*4-HET mice (p_FDR_ = 2.7×10⁻¹¹–0.022; d = -0.66 to -2.29 [95% CI: -2.85, -0.10]), indicating a global effect.

**Figure 3:**
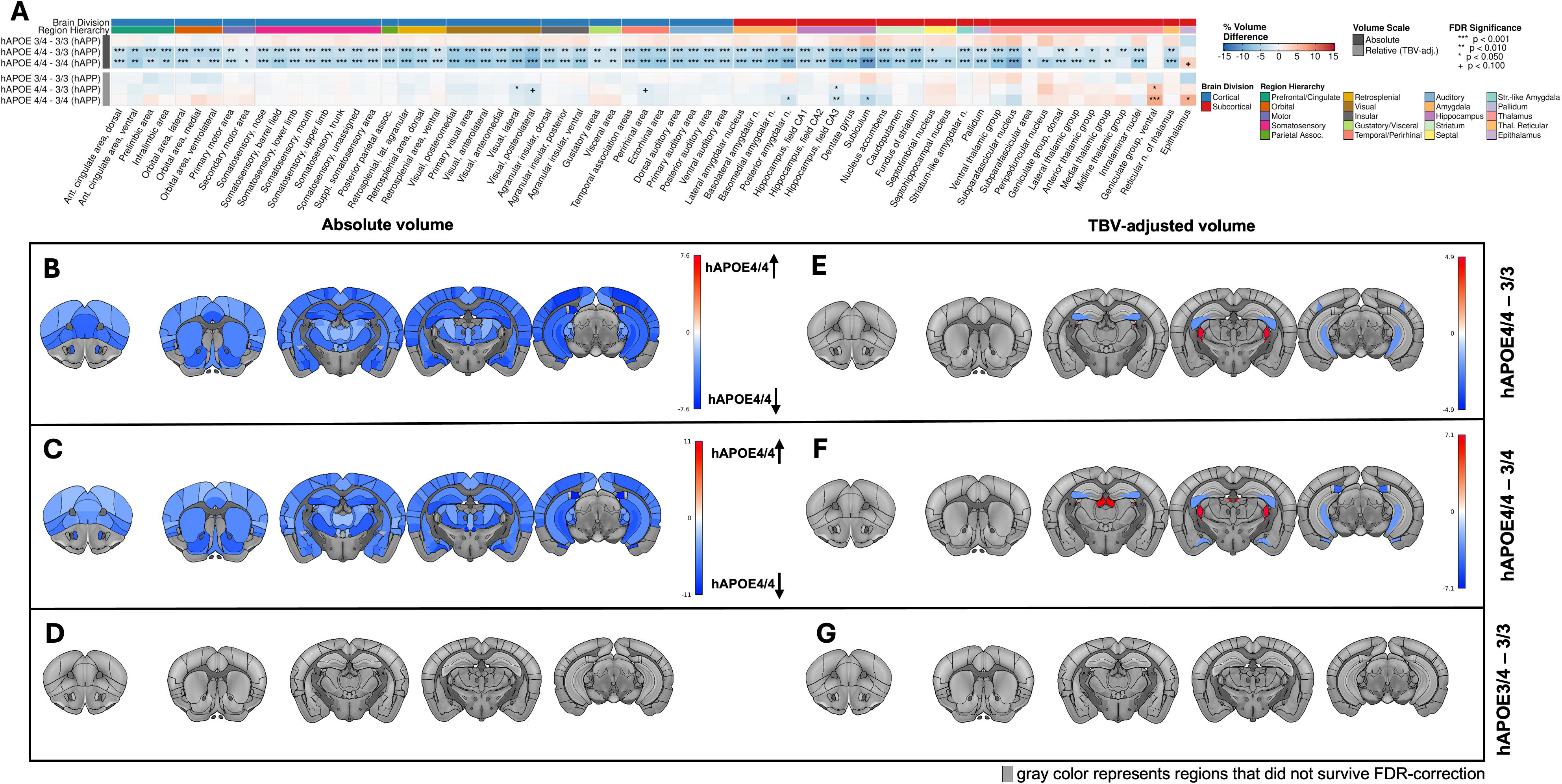
In presence of humanized amyloid precursor protein (APP), two copies of humanized apolipoprotein E4 (h*APOE*4) allele are associated with the lowest brain volume. Results are represented as spatial distribution of regional volume percentage differences (%) linked with the effect of h*APOE*4 carrier status relative to h*APOE*3, on a humanized APP background. All results are FDR-corrected for multiple comparison (p < 0.05). Greater volume is denoted **in red** and lower volume is denoted **in blue.** Regions with no significant differences are presented **in gray.** All results are FDR-corrected for multiple comparison (p < 0.05). **(A) Cumulative region by region difference visualization as a heatmap,** color coded for percentage (%) volume difference, statistical significance and brain region hierarchy. **Left-side** p**anels (B, C, and D)** represent absolute (TBV-unadjusted) volume differences while **right-side panels** (**E, F and G)** represent relative (TBV-adjusted) volume differences.

After TBV adjustment, these global differences were attenuated, and no regionally specific differences emerged between h*APOE*3-HOM and h*APOE*4-HET mice, consistent with the diffuse, non-specific pattern observed in the absolute volumes (**Figure 3D, G**). In contrast, h*APOE*4-HOM mice exhibited a more regionally specific pattern, with smaller volumes localized to visual regions and hippocampal CA3, alongside relative enlargement of the ventral thalamus relative to h*APOE*3-HOM (p_FDR_=0.0206-0.05; d= -0.93 to 1.10 [95% CI: -1.52,1.69]; **Figure 3E**). Additional focal reductions in posterior amygdalar nucleus and subiculum, and enlargement of the epithalamus, were observed in h*APOE*4-HOM relative to h*APOE*4-HET mice (p_FDR_ = 0.0031- 0.0258; d = -1.32 to 1.60 [95% CI: -1.96,2.24]; **Figure 3F**). Collectively, h*APOE*4-HOM drives a global reduction in the *h*APP background that, after normalization, reveals selective regional differences partially mitigated by a single h*APOE*3 allele.

### 2.4. Absolute and TBV-Adjusted Regional Effects of APP Background Stratified by h*APOE* Genotype

Replacing *m*APP with *h*APP resulted in a consistent reduction in regional volumes within each h*APOE* genotype, with a shift in the distribution of effects across genotypes (**Figure 4A-D**). In h*APOE*3-HOM mice, *h*APP effects were heterogeneous, with both larger and smaller regional volumes observed (p_FDR_ = 3.1×10^-11^-0.0302; d=1.2–3.3 [95% CI: -3.28,2.25]; **Figure 4B**). This pattern shifted with increasing h*APOE*4 dose, such that h*APOE*4-HET mice showed predominantly smaller volumes (p_FDR_=1.2×10^-11^-0.0437; d=1.5-4.3 [95% CI: -4.31, -0.11]; **Figure 4C**), and h*APOE*4-HOM mice exhibited a near-uniform reduction across regions (p_FDR_ = 1.6×10^-23^-0.0206; d=1.5–5.1 [95% CI: -5.09,1.49]; **Figure 4D**). Across *h*APOE genotypes, *h*APP mice consistently exhibited larger epithalamic volumes (**Figure 4A**).

**Figure 4:**
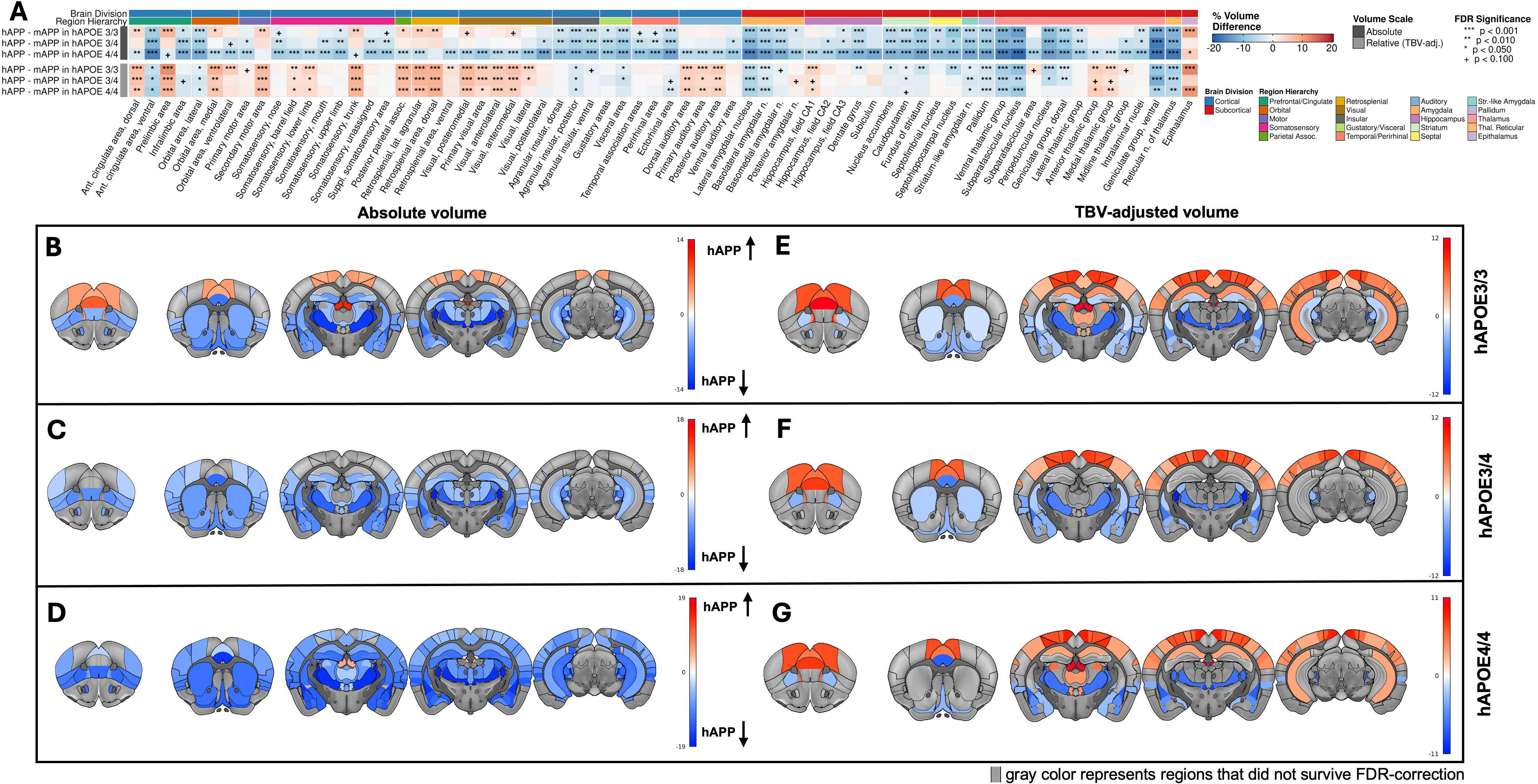
The presence of humanized amyloid precursor protein (*h*APP) across different humanized apolipoprotein (h*APOE*) genotypic combination results in smaller brains with greater representational impact on brain regions involved in cognition. Results shown as spatial distribution of regional volume percentage differences (%) linked with the effect of *h*APP inclusion across each genotypic combination. Greater volume is denoted **in red** and lower volume is denoted **in blue**. Regions with no significant differences are presented **in gray.** All results are FDR-corrected for multiple comparison (p < 0.05). **(A) Cumulative region-by-region difference visualization as a heatmap**, color coded for percentage difference (%), statistical significance and brain region hierarchy. **Left-side panels (B, C, and D)** represent absolute (TBV-unadjusted) volume differences while **right-side panels (E, F and G)** represent relative (TBV-adjusted) volume differences.

After accounting for the globally smaller brains in the *h*APP mice (**Figure 1**), a consistent regional pattern emerged (**Figure 4E-G**). Within h*APOE*4 genotype, *h*APP mice showed relatively greater cortical volumes alongside persistently smaller subcortical volumes, with this cortical–subcortical pattern most clearly defined in h*APOE*4-HOM mice (p_FDR_ = 2.4×10^-12^-0.0487; d=1.2–3.6 [95% CI: -3.58,3.42]; **Figure 4G**). Notable exceptions included larger CA1 volumes, mixed thalamic effects, and persistent epithalamic enlargement across genotypes.

### 2.5. Interaction of humanized APP and humanized *APOE* on regional volume

Difference-in-slope analyses were conducted to assess whether the magnitude of *h*APP-related volume reductions differed by h*APOE* genotype (**Figure 5**). These contrasts indicated that the reduction associated with *h*APP, relative to *m*APP, was consistently greater in h*APOE*4-HOM mice than in h*APOE*3-HOM or h*APOE*4-HET mice, spanning the majority of regions (p_FDR_ = 6.0×10^-6^– 0.0463; d=1.0-3.4 [95% CI: -3.37, -0.09]; **Figure 5A–C**). In contrast, differences between h*APOE*3-HOM and h*APOE*4-HET mice were not observed (all p_FDR_ > 0.05; **Figure 5D**), reflecting their similar volumetric profiles within the *h*APP background.

**Figure 5:**
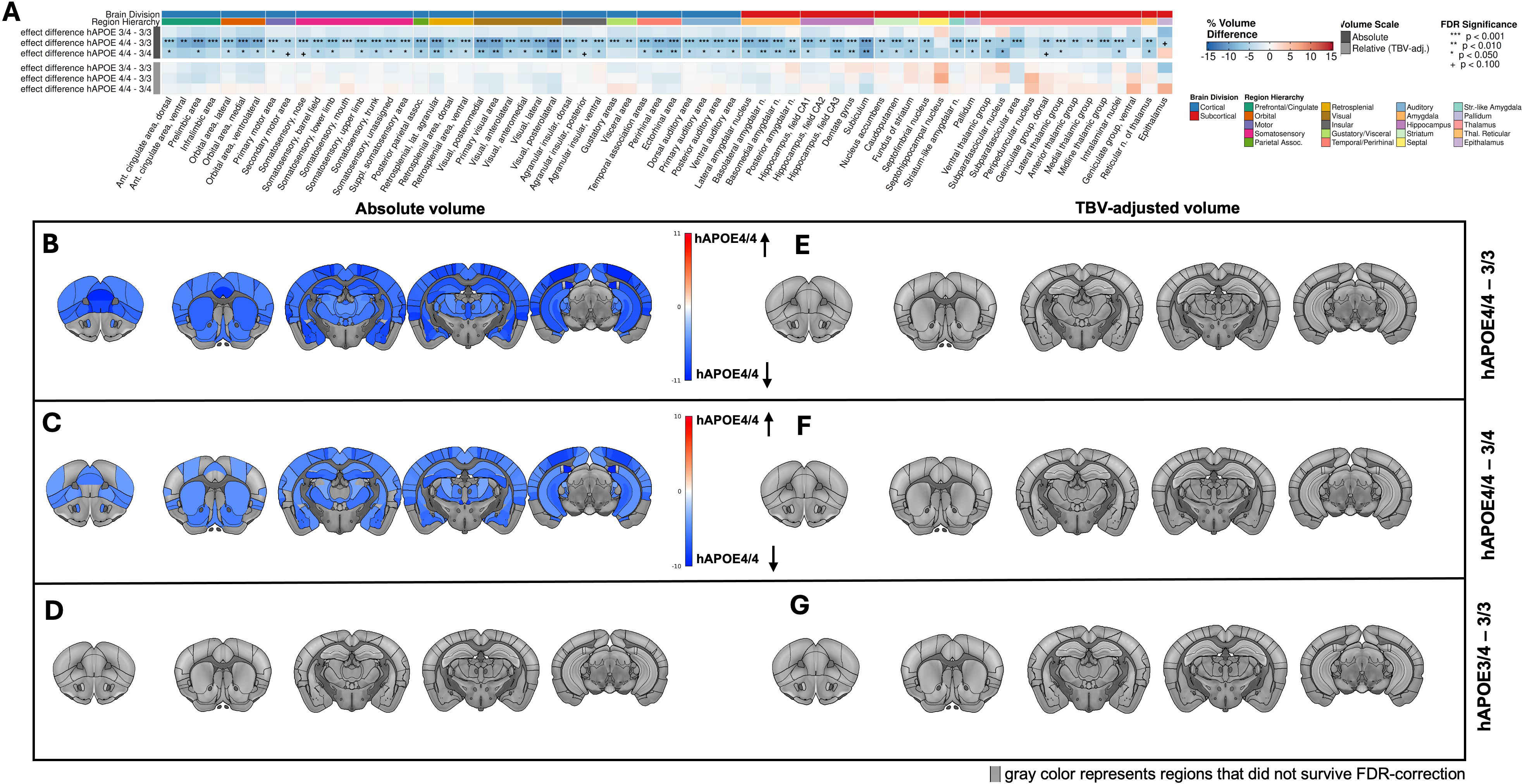
Humanized amyloid precursor protein (*h*APP)-associated reduction across different humanized apolipoprotein (h*APOE*) genotypic combinations showed greatest reductions in the hAPP/h*APOE*4-HOM carriers. Results shown as spatial distribution of regional volume percentage differences (%) linked with the effect of interaction of humanized APP and humanized *APOE*. Lower volume is denoted **in blue.** Regions with no significant differences are presented **in gray.** All results are FDR-corrected for multiple comparison (p < 0.05). **(A) Cumulative region-by-region difference visualization as a heatmap,** color coded for percentage (%) volume difference, statistical significance and brain region hierarchy. **Left-side** p**anels (B, C and D) represent** absolute (TBV-unadjusted) volume differences while **right-side panels** (**E, F and G)** represent relative (TBV-adjusted) volume differences.

After TBV adjustment, differences in *h*APP effect magnitude were attenuated and did not exhibit regionally specific patterns (all p_FDR_ > 0.05; **Figure 5E–G**). However, the residual pattern was directionally consistent with prior observations, with h*APOE*4 carrier mice showing relatively greater cortical reductions and comparatively smaller subcortical reductions than h*APOE*3-HOM mice. *h*APOE4 therefore primarily modulates the magnitude of *h*APP-associated volumetric effects, with limited regional specificity after normalization.

### 2.6. Sex differences

Independent of APP type or *APOE* genotype, female mice had greater TBV than males (mean difference = 1.75%; d = 0.59 [95% CI: 0.26, 0.92]; p = 0.0005). Regionally, this was reflected by greater absolute volumes in females across most of the cortex and subcortex, with no regions showing larger volumes in males (p_FDR_ = 3.0×10^-8^–0.0394; **Figure 6A–B**).

**Figure 6:**
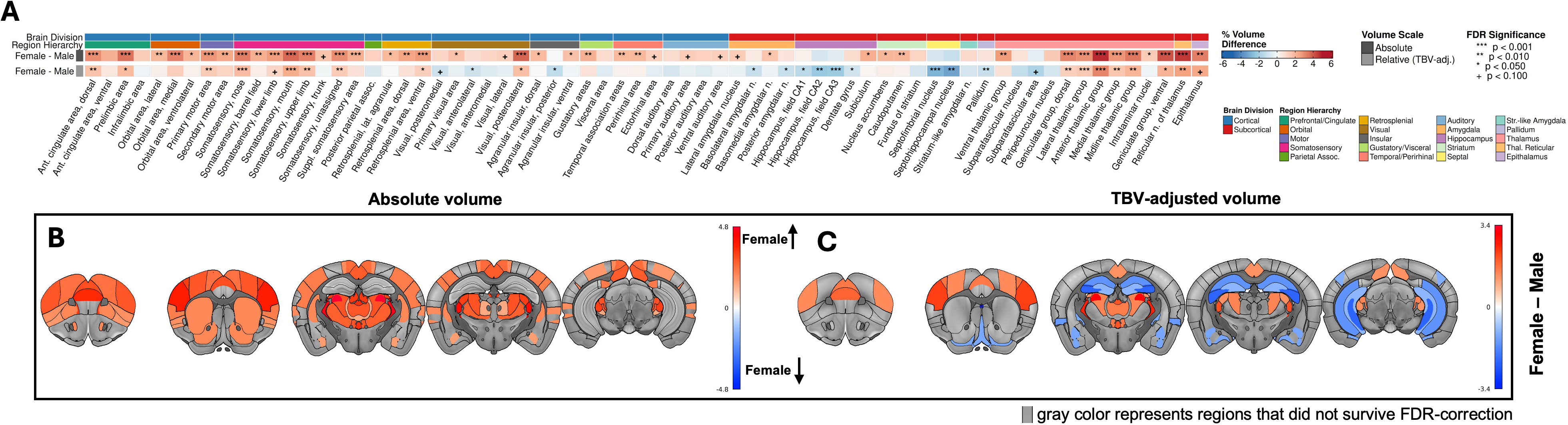
Females had larger brain volume and cortical regions than males, after adjusting for amyloid precursor protein (APP) type and humanized apolipoprotein (h*APOE*) genotype. Results shown as spatial distribution of regional volume differences (%) linked with the effect of sex. Greater volume is denoted **in red** and lower volume is denoted **in blue.** Regions with no significant differences are presented **in gray.** All results are FDR-corrected for multiple comparison (p < 0.05). **A) Region-by-region difference heatmap,** color coded for percentage (%) volume difference, statistical significance and brain region hierarchy. **Left-side panel** showing absolute volumes and **C) Right-side panel** adjusted for differences in total brain volume (TBV).

After TBV adjustment, a regionally specific pattern was evident (**Figure 6C**). Females exhibited relatively greater cortical and thalamic volumes (p_FDR_ = 2.6×10^-6^–0.0191; d = 0.63–1.35 [95% CI: 0.06,1.35]), whereas males showed relatively greater volumes in the hippocampus, amygdala, and striatum (p_FDR_ = 5.3×10^-6^–0.0424; d = 0.55–1.30 [95% CI: 0.06,1.30]). Together, these findings are consistent with sex differences primarily reflecting global scaling of brain volume, with additional region-specific variation after normalization.

### 2.7. Association between behavior and regional brain volumes

Exploratory analyses identified six regions in which associations between regional volume and open field freezing time differed between *h*APP/h*APOE*3-HOM and *h*APP/h*APOE*4-HOM mice. Greater volumes in the striatum-like amygdalar nuclei, posterior agranular insular area, hippocampal CA2, and pallidum were associated with shorter freezing times in *h*APP/h*APOE*4- HOM mice but longer freezing times in *h*APP/h*APOE*3-HOM mice (p_FDR_ = 0.0502 – 0.096; **Supplemental Figure 1**). These associations did not meet a strict p_FDR_ < 0.05 threshold but were retained for descriptive interpretation given their consistency across regions. No significant volumetric associations were observed for novel object recognition discrimination index.

## 3. DISCUSSION

Age, *APOE*4 genotype, and biological sex are major risk factors for late-onset Alzheimer’s disease. The cross-sectional mouse brain volume findings of our prior multi-age study [21] and the aged *m*APP/h*APOE* subset reported herein demonstrate that humanized *APOE*4 does not induce a LOAD-like neuroimaging phenotype, even in aged mice (∼70 years human aged equivalent). h*APOE*4-associated differences in the *m*APP background were small, diffuse, and non-specific, and did not include a pattern of decreased cortical volume or ventricular enlargement [21]. These findings are consistent with vulnerable aging, defined as aging in the presence of neurodegenerative risk factors without confirmed histopathological disease [21].

The present findings extend this, demonstrating that humanized APP reduces total mouse brain volume by approximately 3–10% compared to the *m*APP background, with the greatest reductions observed in *h*APP/h*APOE*4-HOM mice. These volumetric reductions were global in nature, suggesting that *h*APP affects the whole brain rather than the characteristic regional susceptibility observed in LOAD. Interestingly, the effect of h*APOE*4 depended on the APP background, with h*APOE*4-associated volume differences in the *m*APP background differing from those observed in the *h*APP background. Together, these findings suggest an APP– *APOE*4 interaction in which each humanized gene amplifies the structural consequences of the other. However, the combination of these two humanized genes was insufficient to reproduce a LOAD-like volumetric phenotype in these mice.

The ventricular findings further distinguish global volumetric scaling from an AD-like atrophy pattern. *h*APP mice had smaller absolute ventricular volumes than *m*APP mice, with the greatest reduction observed in *h*APP/h*APOE*4-HOM animals. Ventricular volume was reduced by approximately 12.6% in *h*APP/h*APOE*4-HOM relative to *m*APP/h*APOE*4-HOM, and the genotype difference was not significant after total brain volume adjustment. Ventricles therefore scaled with the rest of the brain rather than expanding as if tissue volume had been lost. This pattern is inconsistent with the tissue-loss and CSF-expansion signature of established LOAD. Similarly, relative cortical preservation after adjustment for total brain volume does not reproduce the prominent cortical atrophy observed in established LOAD [6, 34]. The regional overlap with structures affected during LOAD, including thalamic, amygdalar, and hippocampal regions [34–36], indicates translational relevance; however, these differences in the absence of evidence for gray matter atrophy cannot establish a neurodegenerative process.

These findings do not necessarily indicate that the *m*APP/h*APOE* or *h*APP/h*APOE* mouse models are uninformative. Humanized *APOE*4 is associated with endocrine and aging related transcriptomic, cellular, and molecular effects in this model, including inflammation, synaptic biology, cellular metabolism, heterogeneous aging profiles, and endocrine aging predictive of human female LOAD risk [18–20, 37–41]. The present data indicate that these genetically driven risk-related effects can exist, even earlier in life, without producing the macrostructural imaging signature of LOAD. *h*APP may increase the macrostructural susceptibility by converting the h*APOE*4-associated volumetric response from a subtle or diffuse increase into a global decrease. The *h*APP/h*APOE*4-HOM mice specifically capture an APP-dependent consequence of combined genetic risk that is absent in *m*APP/h*APOE*4-HOM mice.

Sex differences further illustrate the distinction between disease risk factors and neurodegenerative features. After accounting for APP and APOE effects, female mice had 1.75% greater total brain volume than males, consistent with prior work identifying sex as a major determinant of brain size across age in the h*APOE* model as well as work in C57BL/6J background strain [42, 43]. This volumetric sex effect does not reflect the higher lifetime risk and greater disease burden in women with AD, or the sex-modifying effects of *APOE*4 on clinical and neuropathological outcomes [44, 45]. Female mice did not show h*APOE*4-dependent volume reductions, indicating that sex-dependent brain volume and sex-dependent AD risk likely reflect partially separable biological pathways involving endocrine, immune, and metabolic factors.

Human studies also demonstrate that genetic risk does not operate independently of the broader aging environment. Neither *APOE* genotype nor AD polygenic risk scores are consistent predictors of cognitive decline [46]. Multidomain mechanisms including epigenetic age, glucose metabolism, and neural compensation rather than merely the absence of AD-related risk likely contribute to neurodegeneration [47, 48]. Therefore, humanized risk genes may establish vulnerability without being sufficient to produce disease-like structural change in isolation.

### 3.1. LIMITATIONS AND FUTURE DIRECTIONS

Several limitations to this study should be acknowledged. First, the current study is cross-sectional, uses ex-vivo fixed-tissue imaging, and did not include any histological assessments. Therefore, we cannot differentiate developmental or constitutive volume effects from neurodegeneration. Second, an aged background-strain cohort was not included, limiting complete characterization of the independent and combined effects of *h*APP and h*APOE*. However, *m*APP/h*APOE* total brain volumes were broadly consistent with published values for adult C57BL/6J mice (mAPP/h*APOE*: 499 mm³; C57BL/6J: 535 mm³). Third, fixation time and storage duration were not explicitly modeled and may have introduced unmeasured confounding due to fixation-related effects. However, all brains underwent the same fixation protocol and our prior work demonstrated limited impacts of storage time on brain volumes [21]. Even if these limitations were fully addressable, they would not be expected to transform the present patterns into LOAD-like imaging phenotypes, given global scaling of brain tissue and ventricles, relative cortical preservation after TBV adjustment, and replication of *m*APP findings.

## 4. CONCLUSION

The utility of these models therefore lies in defining how humanized risk factors alter brain biology in the absence of overt neurodegeneration. The *m*APP/h*APOE* model does not produce AD-like macrostructural atrophy, even at advanced ages, and exhibits a heterogeneous trajectory [21]. The addition of *h*APP creates a distinct and measurable volumetric phenotype, and h*APOE*4 homozygosity increases the magnitude of that effect in a manner that is human-like in principle because it reveals genotype-dependent susceptibility. These data show that humanized genes alone are unlikely to be sufficient to shift the brain from a genetic risk-like phenotype toward disease-like neurodegeneration. Identifying the additional biological, environmental, and systemic factors required for that transition remains an important objective for continued model development [17].

## Data availability statement

The datasets generated and/or analyzed during the current study are available from the corresponding author on reasonable request.

