## Supplemental Figure 1 for "Humanized APP is the primary determinant of regional brain volume in humanized APOE knock-in mice: a cross-sectional ex vivo MRI study"

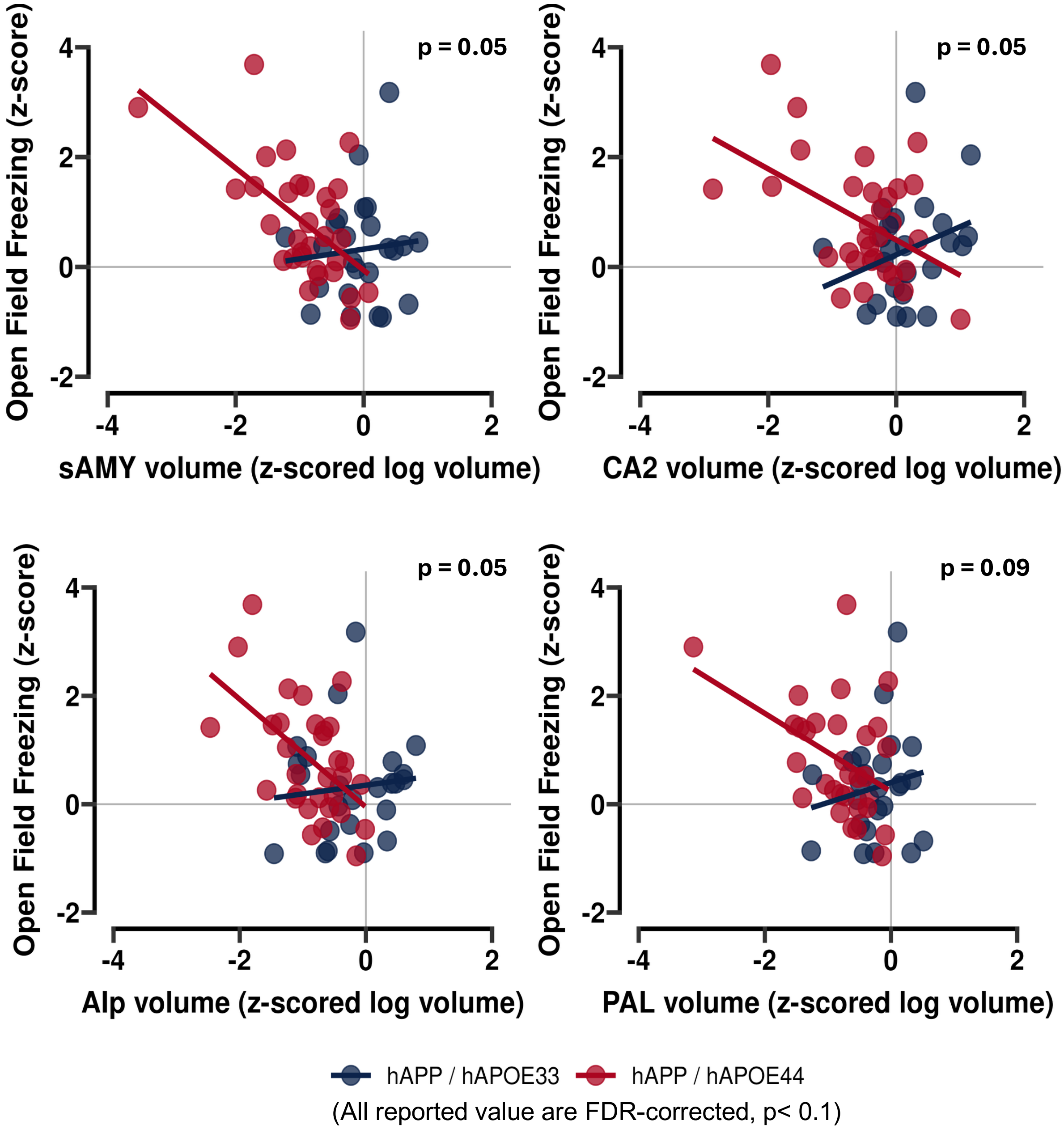


**Supplemental Figure 1: In *h*APP/h*APOE*4-HOM mice smaller regional volumes are associated with greater freezing behavior.** Results are presented as correlation analyses between freezing time and TBV-adjusted regional brain volumes (log scale). All values are z-scored. After controlling for TBV, four regional volumes (in clockwise order from top-left; sAMY – Striatum-like amygdalar nuclei, CA2 of the hippocampus, PAL – Pallidum, and Alp - Agranular insular area) were negatively associated with freezing behavior in *h*APP/h*APOE*4-HOM mice **(in red)** while positively associated with freezing behavior in *h*APP/h*APOE*3-HOM mice **(in blue).** Only FDR-corrected ($p < 0.1$) results are shown.
